# Reduced fetal cerebral blood flow following prenatal drug exposure predicts perinatal mortality

**DOI:** 10.1101/2023.12.01.569643

**Authors:** Siara Kate Rouzer, Anirudh Sreeram, Rajesh Miranda

## Abstract

**Background:** Children exposed prenatally to alcohol or cannabinoids individually can exhibit growth deficits and increased risk for adverse birth outcomes. However, these drugs are often co-consumed and their combined effects on early brain development are virtually unknown. The blood vessels of the fetal brain emerge and mature during the neurogenic period to support nutritional needs of the rapidly growing brain. Teratogenic exposure during this gestational window may therefore impair fetal cerebrovascular development.

**Objective:** To determine whether prenatal polysubstance exposure confers additional risk for impaired fetal-directed blood flow and offspring viability compared to each drug individually.

**Study Design:** We performed high resolution *in vivo* ultrasound imaging in C57Bl/6J pregnant mice. After pregnancy confirmation, dams (*n*=40) were randomly assigned to one of four groups: drug-free control, alcohol-exposed, cannabinoid-exposed or alcohol-and-cannabinoid-exposed. Drug exposure occurred daily between Gestational Days 12-15, equivalent to the transition between the first and second trimesters in humans. Dams first received an intraperitoneal injection of either cannabinoid agonist CP-55940 (750µg/kg) or volume-equivalent vehicle. Then, dams were placed in vapor chambers for 30min of inhalation of either ethanol or room air. Dams underwent ultrasound imaging on three days of pregnancy: Gestational Day 11 (pre-exposure), Gestational Day 13.5 (peri-exposure) and Gestational Day (post-exposure), and were subsequently monitored for health and delivery outcomes.

**Results:** Of all exposure groups, only dams co-exposed to both alcohol and cannabinoids experienced reduced gestational weight gain while undergoing drug treatments. These same co-exposed pregnant mice also demonstrated higher (+42mg/dL) blood ethanol concentrations than dams exposed to alcohol only. All drug exposures decreased fetal cranial blood flow 24-hours after the final exposure episode, though combined alcohol and cannabinoid reduced internal carotid artery blood flow relative to all other exposures. Umbilical artery metrics were not affected by drug exposure, indicating a specific vulnerability of fetal cranial circulation. Cannabinoid exposure significantly reduced cerebroplacental ratios, mirroring prior findings in marijuana-exposed human fetuses. Post-exposure cerebroplacental ratios significantly predicted subsequent perinatal mortality (*p*=0.019, area under the curve, 0.772; sensitivity, 81%; specificity, 85.70%) and retroactively diagnosed prior drug exposure (*p*=0.005; AUC, 0.861; sensitivity, 86.40%; specificity, 66.7%).

**Conclusion(s):** Fetal cerebrovasculature is significantly impaired by exposure to alcohol or cannabinoids, and co-exposure confers additional risk for adverse birth outcomes. Considering the rising potency and global availability of cannabis products, there is an imperative for research to explore translational models of prenatal drug exposure, including polysubstance models, to inform appropriate strategies for treatment and care in pregnancies affected by drug exposure.

**Key Points:** *Question:* Does simultaneous prenatal exposure to alcohol and cannabinoids present significant additional risk to fetal health compared to each drug individually?

*Findings:* Maternal murine ultrasound analyses showed that alcohol and cannabinoid exposure, individually, reduced fetal cerebral arterial blood flow metrics. Notably, polysubstance-exposed fetuses demonstrate the worst cerebral hemodynamics, and reductions in fetal blood flow significantly predict subsequent perinatal offspring mortality.

*Meaning:* Prenatal drug exposure persistently reduces fetal-directed blood flow, which can disrupt normal embryonic growth and neural development, and polysubstance exposure augments deficits specifically in cerebral arterial blood flow.

## Introduction

Co-consumption of alcohol with cannabis is a rapidly emerging public health problem, partly due to widespread marijuana legalization and increased availability of potent synthetic cannabinoids ^1^. Approximately one-third of young adults in the United States report using either drug within the past month ^2^. Moreover, ∼8-12% of infants are prenatally exposed to ethanol ^3–5^, and ∼22-40% are exposed to cannabis during the last month of pregnancy ^6,7^. Importantly, young adults who use both substances preferentially engage in the practice of simultaneous alcohol and cannabinoid (SAC) consumption ^8^, which amplifies each drug’s psychological effects ^9^.

However, the consequences of SAC for pregnancy health and fetal development are currently poorly understood. Individually, prenatal exposures to either alcohol or cannabinoids have similar physical and developmental consequences for exposed offspring, each resulting in growth restriction and reduced birth weights ^10–12^, increased rates of preterm delivery ^10,13–15^, and increased health risks associated with delivery ^13,16^. Cannabis exposure, in particular, has been tied to increased need for neonatal intensive care services ^17^ and greater risk of fetal demise ^16^. Both ethanol and cannabinoids cross the placental barrier ^18,19^, and consequently, have direct effects on the developing fetus. While little is known about the specific effects of co-use, two recent papers in mouse ^20^ and zebrafish ^21^ models indicate that teratogenic effects of prenatal alcohol exposure are amplified by concurrent cannabinoid exposure. Therefore, SAC is likely to result in increased deficits compared to either prenatal alcohol or cannabinoid exposures alone.

Here, we assessed the vulnerability of fetal brain vasculature following maternal SAC. Studies in children with fetal alcohol spectrum disorders, as well as translational animal models, have shown that prenatal alcohol exposure results in structural and functional changes in fetal brain circulation and arterial vasodilation, leading to both acute and persistent loss of cerebral blood flow to the developing brain (for review, see ^22^). To date, few studies have explored the effects of *in utero* cannabinoid exposure on fetal cerebrovasculature. Ultrasound imaging in pregnant women during the second trimester has revealed that daily marijuana use is associated with higher umbilical artery systolic:diastolic ratios when compared to controls ^23^. This anomaly persisted into the third trimester, with marijuana-exposed fetuses demonstrating a higher incidence of growth restriction, and some exhibiting absent or reversed end-diastolic blood flow in the umbilical artery, an outcome that predicts fetal distress ^24^. Moreover, marijuana-exposed fetuses with growth restriction exhibit abnormally low cerebroplacental ratios (CPR), the ratio of middle cerebral artery to umbilical artery blood flow ^25^. Low fetal CPR was also associated with low birth weight and an increased need for neonatal intensive care. Rapid reduction in fetal cerebral vessel diameter, length fraction, and area density have been further observed in murine models following acute maternal exposure to synthetic cannabinoid CP-55940 ^26^.

To our knowledge, no studies have specifically addressed the impact of SAC on fetal vascular function. Therefore, we investigated the effects of singular or simultaneous exposure to alcohol and cannabinoids during a critical period of cerebral cortical plate neurogenesis and angiogenesis ^27,28^) on fetal blood flow and subsequent viability at delivery. Our results indicate that both forms of prenatal drug exposure contribute to persistent reductions in fetal-directed cerebral blood flow, with SAC augmenting certain deficits further. Moreover, SAC was associated with low CPR and the highest rates of perinatal mortality among offspring compared to all other exposure conditions.

## Methods

### Breeding paradigm

All procedures were performed with Texas A&M’s Institutional Animal Care Committee oversight. One male and two female C57Bl/6J female mice (Harlan laboratories, Houston, TX) were temporarily co-housed overnight for mating. Pregnancies were confirmed the next morning (Gestational Day [G]1) by the presence of a sperm plug. Dams were weighed on G1 and G10 to verify weight gain consistent with pregnancy, and subsequently on G15 and G18, to determine effects of drug exposure on gestational weight gain. After pregnancy confirmation, dams were randomly assigned to one of four groups: drug-free controls, only alcohol-exposed (Alcohol), only cannabinoid-exposed (Cannabinoid) or alcohol-and-cannabinoid-exposed (Alcohol+Cannabinoid).

### Prenatal drug exposure paradigm

Exposures occurred daily between G12-15 (Fig. 1B). First, dams received an intraperitoneal injection of either cannabinoid agonist CP-55940 (Tocris Bioscience, Bristol, UK; 750µg/kg dissolved in 10% DMSO/saline), or vehicle (volume-equivalent 10% DMSO/saline). Next, dams were placed in vapor chambers (passive e-vape system, La Jolla Alcohol Research Inc., La Jolla, CA) for 30min of inhalation of either ethanol (95% ethanol) or room air, as previously described ^29^. Following exposure, dams were assessed for exposure-induced loss of righting reflex (LORR) and tail blood was collected (20µL) in heparinized capillary tubes to assess blood alcohol concentrations by gas chromatography, as previously described ^30–33^.

**Figure 1.**
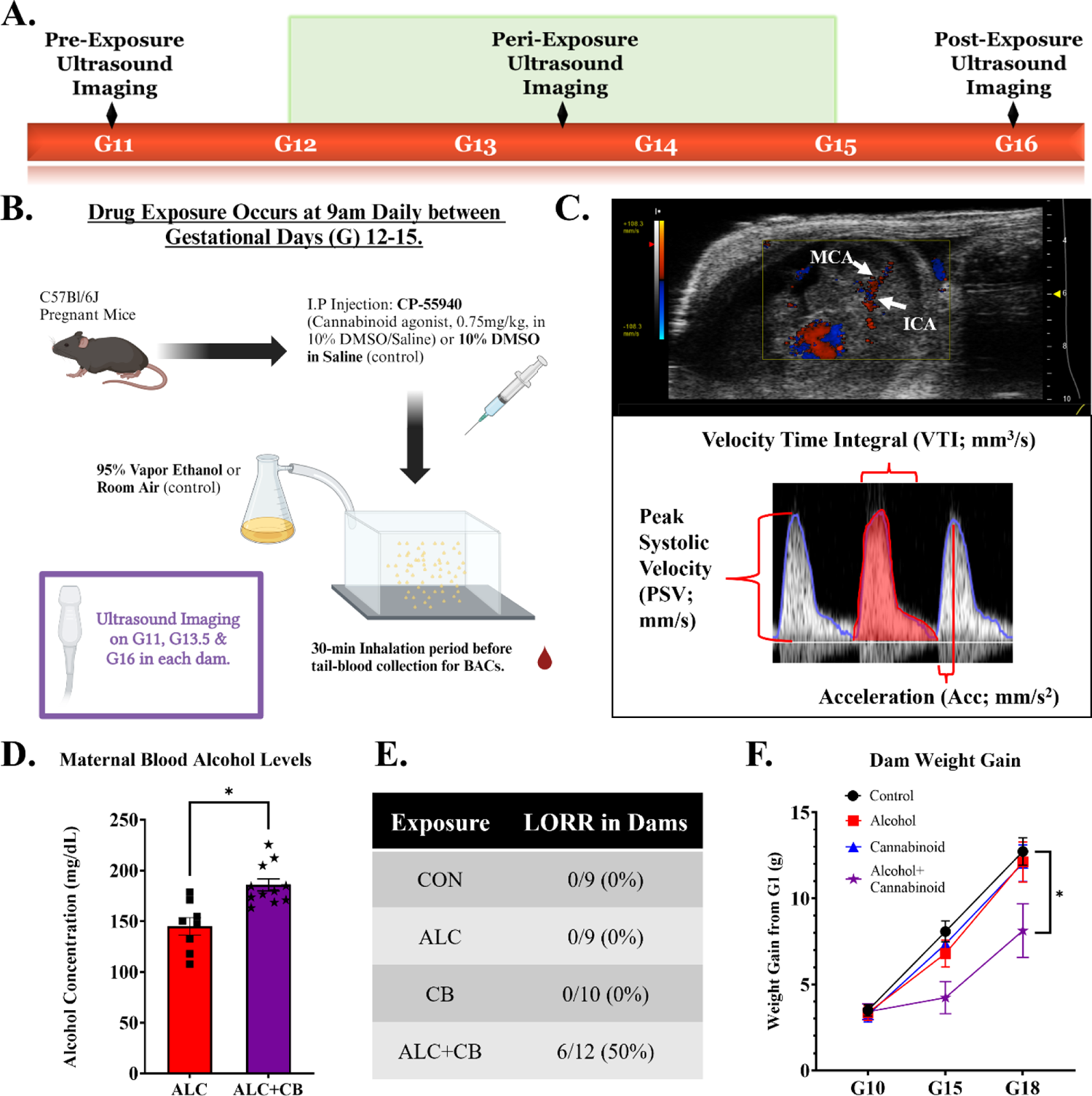
Experimental outline and polysubstance exposure characterization. **1A)** Experimental timeline for prenatal drug exposure and ultrasound imaging data collection. **1B)** Summary figure of the daily prenatal drug exposure procedure. **1C)** Example fetus, G13.5, imaged using Vevo2100, with example waveform analysis figure. **1D)** Average blood alcohol concentrations among alcohol-exposed subjects. **1E)** Rates of loss of righting reflex (LORR) in drug-exposed dams immediately following drug exposure. **1F)** Gestational weight gain across drug-exposed dams. **Abbreviations**: G=Gestational Day, BAC= Blood Alcohol Concentration, MCA=Middle Cerebral Artery, ICA=Internal Carotid Artery, LORR = Loss of Righting Reflex, CON=Control, ALC=Alcohol, CB=Cannabinoid, ALC+CB=Alcohol+Cannabinoid. **Symbols:** * indicates *p* < 0.05.

### Ultrasonography

Each pregnant dam underwent ultrasound imaging at G11 (pre-drug exposure), G13.5 (peri-drug exposure) and G16 (post-drug exposure) (Fig. 1A&B), as we previously described ^34,35^. Briefly, dams were anesthetized with isoflurane (2-3%), and maintained supine on a temperature-controlled sensor platform to monitor maternal electrocardiogram, respiration and core body temperature (Visualsonics, Toronto, Canada). For each pregnant dam, a single fetus located at the base of the uterine horn was selected for repeated imaging. Doppler measurements for umbilical arteries and fetal internal carotid/middle cerebral arteries were obtained using a high-frequency VEVO2100 ultrasound imaging machine coupled to a MS550D Microscan™ transducer with a center frequency of 40MHz (Visualsonics, Canada). Following drug exposure, dams were monitored daily by researchers to assess maternal health outcomes and to verify date of parturition.

### Data analysis and statistics

Ultrasound subjects were assigned random identification codes and recordings were analyzed by an experimenter blinded to the exposure condition. Ultrasound imaging data were analyzed using the VEVO2100 measurement and analysis software (Visualsonics, Ca, Fig. 1C) to assess Velocity-Time Integral (VTI, in mm^3^/sec, a measure of cardiac stroke volume through a specific blood vessel), peak systolic velocity (PSV, the peak quantity of blood flow during a systolic pulse) and Acceleration (Acc, in mm/sec^2^, a measure of arterial resistance) ^34,36,37^. Each data point reflects measurements from one fetus, from one pregnant dam. Based on preliminary data, a power analysis (G*Power 3.1.9.4) for a two-way (gestational day x exposure) analysis of variance (ANOVA) and an alpha of 0.05 suggested a sample size of 7 pregnant dams per group. Results are presented as mean ± SEM for each group.

Gestational weight gain and blood flow metrics were analyzed using a mixed-effects ANOVA, with independent variables of gestational day (within-subjects) and drug exposure. In the event of a significant effect of exposure, or an interaction of gestational day x exposure, post-hoc Tukey tests were performed comparing all exposure groups within-day. To determine if drug exposure produced significant, within-animal changes in blood flow growth metrics, one-sample t-tests were performed within each exposure group. One-way ANOVAs were also used to compare drug-induced changes between exposure groups. Pearson’s *r* correlations were performed between fetal blood flow metrics and maternal/fetal mortality, and Kaplan-Meier Survival Analyses were used to calculate the probability of survival within exposure groups.

Post-hoc pairwise comparisons then determined exposure differences in survival. Receiver operating characteristic (ROC) analyses were performed to assess the accuracy of blood flow metrics in predicting either future fetal mortality or prior drug exposure. Gestational blood alcohol concentrations were assessed using a standard t-test. All data were assessed for outliers using the ROUT method of regression with a false-discovery rate of 1%, and identified outliers were removed from statistical analyses. Group differences were considered significant at *p*≤0.05. All statistics were performed using GraphPad Prism 9, except for ROC analyses (SPSS v28, IBM).

## Results

### SAC increased blood alcohol concentrations and LORR, and reduced gestational weight gain

A total of 40 dams were exposed in our paradigm (Control: 9, Alcohol: 9, Cannabinoid: 10, Alcohol+Cannabinoid: 12). Blood alcohol concentrations achieved in alcohol-exposed pregnancies were significantly lower (∼22%) than concentrations achieved following combined exposure to alcohol and CP-55940 [Fig. 1D; *t*(6)=3.090, *p*=0.021], despite identical alcohol exposure conditions. Moreover, only Alcohol+Cannabinoid dams experienced motor impairment, measured by LORR (Fig. 1E). All pregnant animals experienced gestational weight gain (significant main effect of gestational day [*F*(2,50)=238.4, *p*<0.001]), with no group differences in weight gain on G10, prior to exposure (eTable 1). However, there was a significant interaction between gestation day x exposure [*F*=(6,64)=5.146, *p*<0.001], due to decreased weight gain in Alcohol+Cannabinoid dams at the end of the exposure period (G15, *p*=0.014 compared to control dams, Fig. 1F).

From each exposure group, 7-8 dams were selected to undergo ultrasound imaging prior to (G11), during (G13.5) and after (G16) the drug exposure period. Indices for VTI, Acc and PSV were computed for umbilical and fetal middle cerebral and internal carotid arteries. Blood flow metric effect sizes are listed in Table 1. Importantly, no exposure condition significantly changed blood flow metrics in the umbilical artery (eFig. 1).

**Table 1.** Effects of prenatal drug exposure on fetal blood flow metrics & associated effect sizes. Table of blood flow metric comparisons & associated effect sizes (Hedge’s *g*, reported as Effect Size [95% CI upper, lower limits]). Bold text indicates accompanying *p*-value < 0.05. Italicized text indicates accompanying *p*-value < 0.1. **Abbreviations:** CON=Control, ALC=Alcohol, CB=Cannabinoid, ALC+CB= Alcohol+Cannabinoid, VTI=Velocity-Time Integral, MCA=Middle Cerebral Artery, ICA=Internal Carotid Artery, UA=Umbilical Artery.

| <u>VTI</u> |  | Main effect:<br>exposure | Interaction:<br>exposure x day | Hedge's g: CON vs<br>Alcohol (ALC) | Hedge's g: CON vs<br>Cannabinoid (CB) | Hedge's g: CON<br>vs ALC+CB |
| --- | --- | --- | --- | --- | --- | --- |
|  |  | *p-value* | *p-value* |  |  |  |
| **MCA** | G11 | **0.0207** | **0.0006** | 0.078 [-1.715, 1.871] | 0.628 [-0.411, 1.667] | 0.49 [-0.54, 1.519] |
|  | G13.5 |  |  | 1.195 [0.094, 2.295] | 1.279 [0.166, 2.391] | 0.717 [-0.33, 1.763] |
|  | G16 |  |  | **1.374 [0.247, 2.502]** | **1.267 [0.156, 2.378]** | **1.735 [0.546, 2.924]** |
| **ICA** | G11 | **0.0022** | **0.0306** | 0.448 [-0.579, 1.475] | 0.065 [-0.95, 1.079] | 0.176 [-0.84, 1.192] |
|  | G13.5 |  |  | 1.209 [0.106, 2.311] | 0.22 [-0.797, 1.237] | 0.69 [-0.354, 1.734] |
|  | G16 |  |  | 0.84 [-0.218, 1.897] | 0.692 [-0.352, 1.736] | **1.919 [0.694, 3.144]** |
| **UA** | G11 | 0.6007 | 0.7432 | 0.121 [-0.894, 1.136] | 0.22 [-0.797, 1.238] | 0.186 [-0.83, 1.203] |
|  | G13.5 |  |  | 0.897 [-0.167, 1.961] | 0.812 [-0.243, 1.867] | 0.829 [-0.227, 1.886] |
|  | G16 |  |  | 0.514 [-0.516, 1.545] | 0.23 [-0.788, 1.248] | 0.592 [-0.444, 1.629] |
| <u>Acceleration</u> |  | Main effect:<br>exposure | Interaction:<br>exposure x day | Hedge's g: CON vs<br>Alcohol (ALC) | Hedge's g: CON vs<br>Cannabinoid (CB) | Hedge's g: CON<br>vs ALC+CB |
|  |  | *p-value* | *p-value* |  |  |  |
| **MCA** | G11 | **0.0011** | 0.0994 | 0.267 [-0.441, 0.975] | 0.193 [-0.51, 0.895] | 0.006 [-0.701, 0.714] |
|  | G13.5 |  |  | 1.114 [0.024, 2.204] | 0.961 [0.239, 1.682] | 1.615 [0.853, 2.377] |
|  | G16 |  |  | **1.402 [0.271, 2.534]** | **1.447 [0.733, 2.162]** | **1.319 [0.612, 2.026]** |
| **ICA** | G11 | 0.0874 | 0.3412 | 0.577 [0.408, 0.746] | 0.179 [-0.521, 0.878] | 0.337 [-0.375, 1.048] |
|  | G13.5 |  |  | 0.629 [0.431, 0.828] | 0.452 [-0.257, 1.161] | 0.427 [-0.276, 1.129] |
|  | G16 |  |  | 0.347 [-0.353, 1.047] | 0.713 [0.012, 1.415] | **1.134 [0.432, 1.836]** |
| **UA** | G11 | 0.5798 | 0.8334 | 0.205 [-0.542, 0.951] | 0.101 [-0.914, 1.116] | 0.148 [-0.868, 1.164] |
|  | G13.5 |  |  | 0.968 [0.21, 1.727] | 0.415 [-0.61, 1.44] | 1.756 [0.563, 2.949] |
|  | G16 |  |  | 0.033 [-0.715, 0.78] | 0.384 [-0.639, 1.408] | 0.355 [-0.667, 1.377] |
| <u>Peak Systolic</u><br><u>Velocity</u> |  | Main effect:<br>exposure | Interaction:<br>exposure x day | Hedge's g: CON vs<br>Alcohol (ALC) | Hedge's g: CON vs<br>Cannabinoid (CB) | Hedge's g: CON<br>vs ALC+CB |
|  |  | *p-value* | *p-value* |  |  |  |
| **MCA** | G11 | **0.0437** | **0.011** | 0.044 [-0.879, 0.967] | 0.454 [-0.423, 1.331] | 0.431 [-0.442, 1.304] |
|  | G13.5 |  |  | 0.837 [-0.221, 1.895] | 0.779 [-0.252, 1.81] | 0.302 [-0.529, 1.132] |
|  | G16 |  |  | **1.189 [0.089, 2.289]** | **1.247 [0.339, 2.155]** | **1.277 [0.421, 2.132]** |
| **ICA** | G11 | **0.0263** | **0.0326** | 0.103 [-0.737, 0.942] | 0.823 [-0.057, 1.704] | 0.513 [-0.458, 1.483] |
|  | G13.5 |  |  | 1.209 [0.23, 2.188] | 0.487 [-0.627, 1.601] | 0.615 [-0.24, 1.469] |
|  | G16 |  |  | 0.766 [-0.07, 1.602] | 0.652 [-0.132, 1.437] | **1.601 [0.751, 2.451]** |
| **UA** | G11 | 0.5708 | 0.5999 | 0.232 [-0.603, 1.067] | 0.294 [-0.726, 1.313] | 0.403 [-0.621, 1.428] |
|  | G13.5 |  |  | 0.888 [-0.074, 1.85] | 0.7 [-0.345, 1.745] | 1.158 [0.062, 2.253] |
|  | G16 |  |  | 0.445 [-0.401, 1.29] | 0.342 [-0.679, 1.364] | 0.58 [-0.455, 1.616] |

### All drug exposures significantly and persistently reduced middle cerebral artery blood flow metrics

All drug exposures - Alcohol, Cannabinoid & Alcohol+Cannabinoid - reduced fetal middle cerebral artery VTI relative to controls, but only after the exposure period, i.e., at G16 [significant interaction between gestational day and exposure: *F*(6,50)=4.853, *p*<0.001, Figs. 2A&B, eTable 2). VTI significantly increased in control fetuses between G11 and G16 (*p*=0.002). This age-related increase was partly attenuated in Alcohol fetuses (*p*=0.049) and completely abolished in Cannabinoid and Alcohol+Cannabinoid fetuses (all *p*’s >0.357; Fig. 2C; eTables 3&4). As VTI measurements can reflect changes in arterial resistance and/or the quantity of blood flow, we also assessed middle cerebral artery Acc and PSV (eFig. 2). Like VTI, all forms of prenatal drug exposure significantly reduced Acc (all *p*’s < 0.005; eTable 5) and PSV (all *p*’s < 0.005; eTable 6) relative to controls on G16, though drug-exposed groups did not differ statistically from each other.

**Figure 2.**
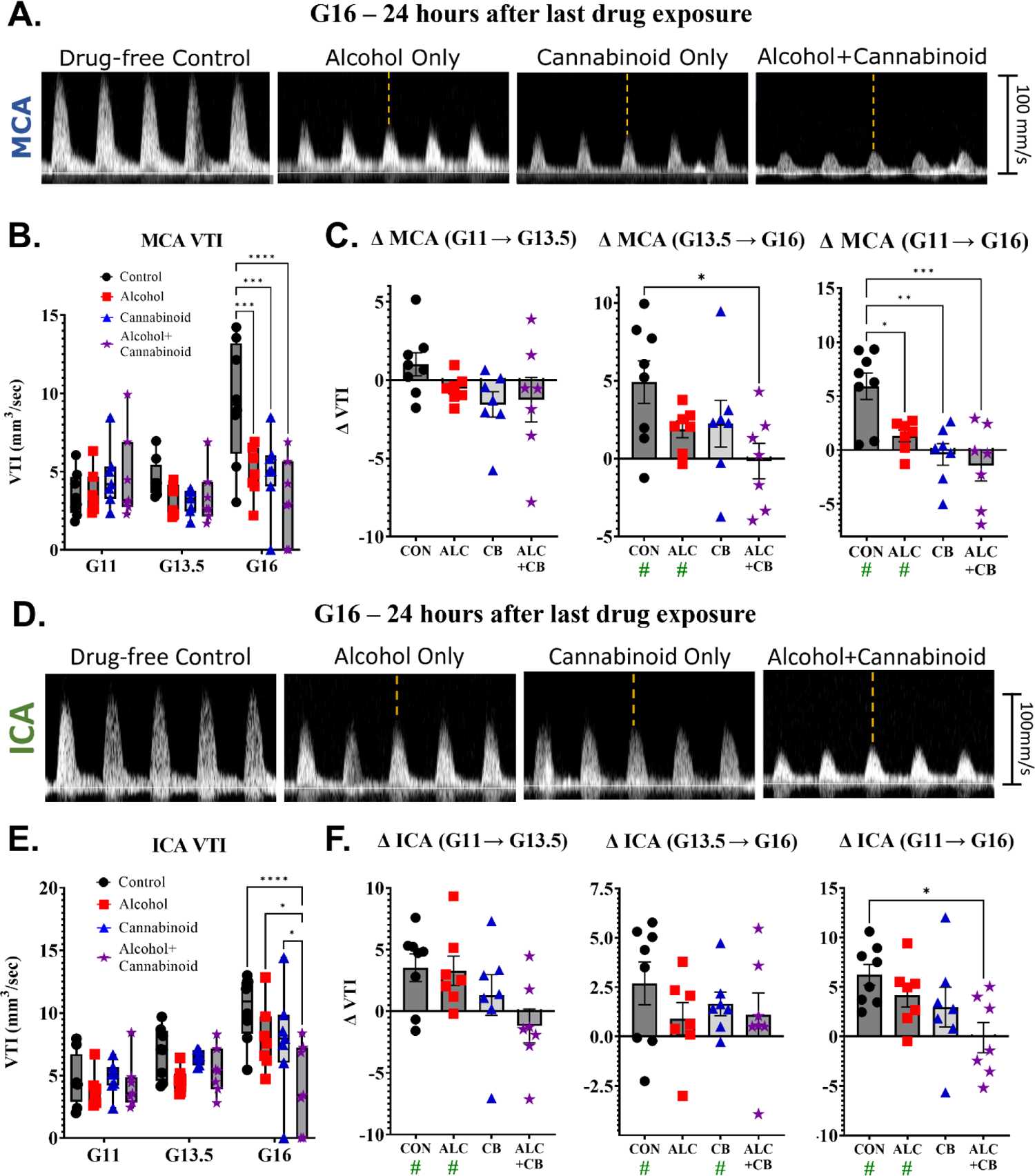
Drug-associated changes in fetal blood flow in the middle cerebral and internal carotid arteries. **2A)** Representative examples of systolic pulse waveforms from the fetal middle cerebral artery across drug groups on Gestational Day 16. **2B)** Velocity-Time Integral measurements from the fetal middle cerebral artery across drug groups and assessment days. **2C)** Within-fetus changes in middle cerebral artery Velocity-Time Integral measurements across drug groups and assessment days. **2D)** Representative examples of systolic pulse waveforms from the fetal internal carotid artery across drug groups on Gestational Day 16. **2E)** Velocity-Time Integral measurements from the fetal internal carotid artery across drug groups and assessment days. **2F)** Within-fetus changes in internal carotid artery Velocity-Time Integral measurements across drug groups and assessment days. **Abbreviations**: G=Gestational Day, MCA=Middle Cerebral Artery, ICA=Internal Carotid Artery, VTI=Velocity-Time Integral, CON=Control, ALC=Alcohol, CB=Cannabinoid, ALC+CB=Alcohol+Cannabinoid. **Symbols:** Green # indicates a statistically significant change from 0. * indicates *p* < 0.05, ** indicates *p* < 0.01, *** indicates *p* < 0.001, **** indicates *p* < 0.0001.

### SAC distinctly reduced internal carotid artery metrics on G16 compared to single-drug exposure groups

As with the middle cerebral artery, drug exposures reduced VTI in the fetal internal carotid artery only after the exposure period [significant interaction between gestational day and exposure: *F*(6,75)=2.478, *p*=0.031]. However, post-hoc analyses revealed that this effect was driven by the Alcohol+Cannabinoid group, which significantly reduced VTI relative to all other exposures (Control, Alcohol & Cannabinoid: all *p*’s < 0.027). Neither Alcohol nor Cannabinoid fetuses differed from controls (*p*’s>0.150; Figs. 2D&E, eTable 7). While VTI increased in control fetuses from G11 to G16 (*p*<0.001), this age-related increase was attenuated in Alcohol (*p*=0.032) and Cannabinoid (*p*=0.032) fetuses (Fig. 2F), and abolished in Alcohol+Cannabinoid fetuses (*p*=0.939). This VTI growth significantly differed between Control and Alcohol+Cannabinoid fetuses exclusively (*p*=0.022; eTable 8). Alcohol+Cannabinoid fetuses were also the only group to show significant reductions in internal carotid artery Acc on G16 (*p*=0.015; eFig. 3A) relative to controls (eTable 9). Furthermore, on G16, Alcohol+Cannabinoid fetuses demonstrated reduced PSV compared to all other exposure groups (Control, Alcohol & Cannabinoid: all *p*’s < 0.059; eFig. 3B), with no other group differences observed (eTable 10).

**Figure 3.**
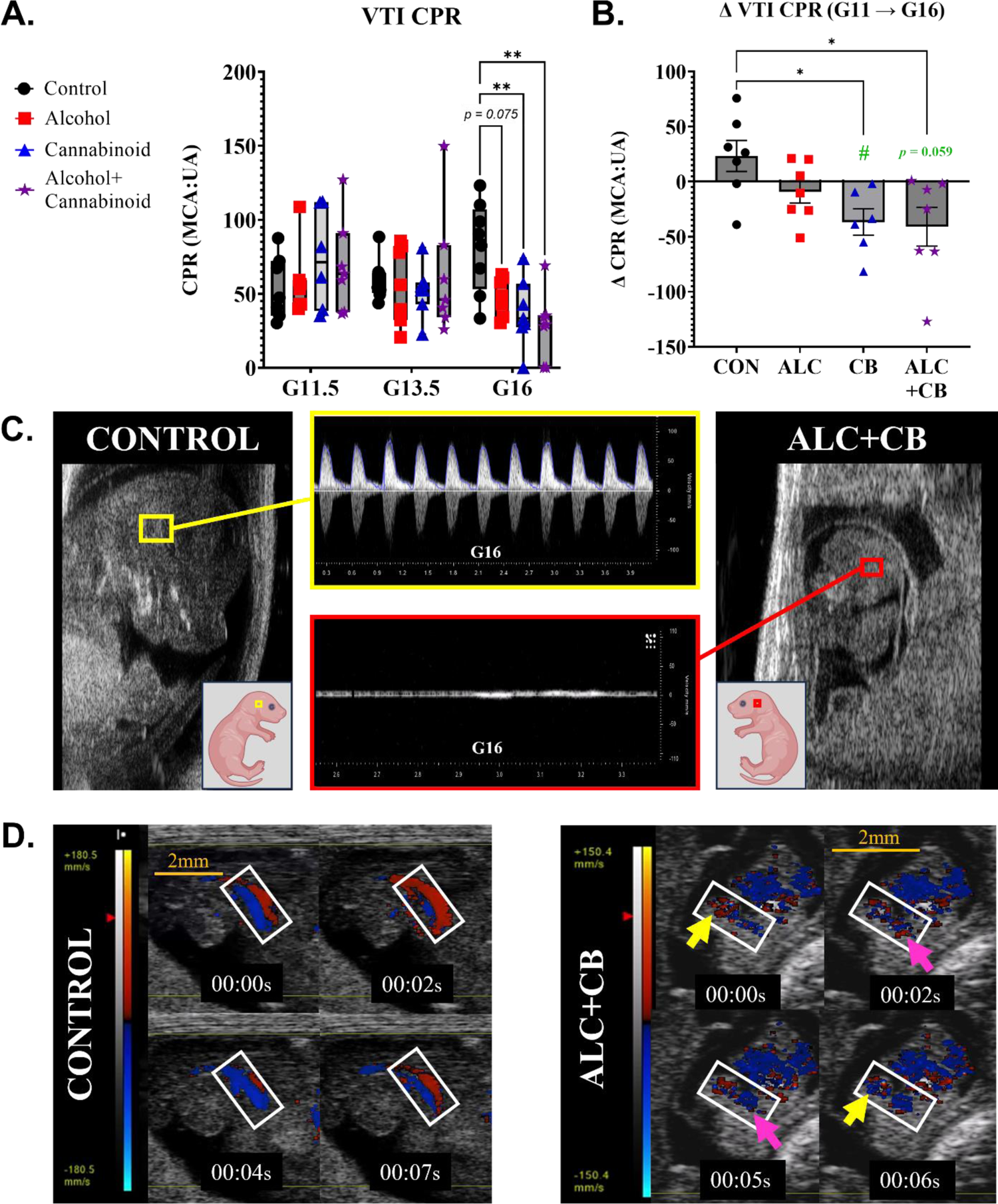
Changes in fetal cerebroplacental ratios following prenatal drug exposure. **3A)** Cerebroplacental ratios calculated within-animal across drug groups and assessment days. CPR = [MCA VTI/ UA VTI] * 100. **3B)** Change within-animal in cerebroplacental ratios prior to drug exposure (Gestational Day 11) and following drug exposure (Gestational Day 16). **3C)** Gestational Day 16 images of a fetus with no drug exposure (left, yellow) and a growth-restricted fetus with alcohol+cannabinoid exposure (right, red). Waveform images demonstrate a lack of directional blood flow in alcohol+cannabinoid fetus. **3D)** Series of Color doppler images of a drug-free control fetus (left) and a growth-restricted alcohol+cannabinoid exposed fetus (right). In the control fetus, the umbilical artery and vein run parallel to each other without overlapping. In contrast, within the alcohol+cannabinoid exposed fetus, red and blue colors overlay within the same regions (arrows), indicating a loss of directional blood flow in the umbilical artery. **Abbreviations**: G=Gestational Day, VTI=Velocity-Time Integral, CPR=Cerebroplacental Ratio, CON=Control, ALC=Alcohol, CB=Cannabinoid, ALC+CB=Alcohol+Cannabinoid. **Symbols:** Green # indicates a statistically significant change from 0. * indicates *p* < 0.05, ** indicates *p* < 0.01.

### Prenatal cannabinoid exposure diverts fetal blood away from the brain

Prior investigations in human fetuses link marijuana exposure to reduced CPR ^25^. Here, assessments of CPR for mouse fetuses, as a ratio-percent of VTI in the middle cerebral artery relative to the umbilical artery, revealed a significant interaction between gestational day and exposure [*F*(6,48)=3.082, *p*=0.012]. Post-hoc comparisons (Fig. 3A) showed that, relative to controls, CPR declined on G16 in both cannabinoid-exposed groups: Cannabinoid (*p*=0.009) and Alcohol+Cannabinoid (*p*=0.001) (eTable 11). Similarly, within-animal, only cannabinoid-exposed dams demonstrated reduced CPR following the drug exposure period (Fig. 3B, eTable 12). Reduced CPR in prior investigations of marijuana-exposed human fetuses was also accompanied, in some cases, by reversed end-diastolic blood flow ^23^, an index of fetal distress ^24^. Similar outcomes were observed in a few growth-restricted, cannabinoid-exposed fetal mice (1 Cannabinoid fetus, 2 Alcohol+Cannabinoid fetuses) assessed on G16, including near loss of blood flow in the middle cerebral artery (Fig. 3C, with an age-matched Control fetus for comparison; Videos 1&2) and flow-reversal in umbilical circulation (Fig. 3D; Videos 3&4).

### Fetal blood flow metrics predict drug-exposure-associated perinatal fetal mortality

Kaplan-Meier analysis showed that drug-exposed groups, but not controls, experienced compromised fetal survival [Fig. 4A; X^2^(3, N=40)=15.33, *p*=0.002]. Log-Rank (Mantel-Cox) analyses revealed that Control and Alcohol fetuses did not differ in survival rates (*p*=0.171).

**Figure 4.**
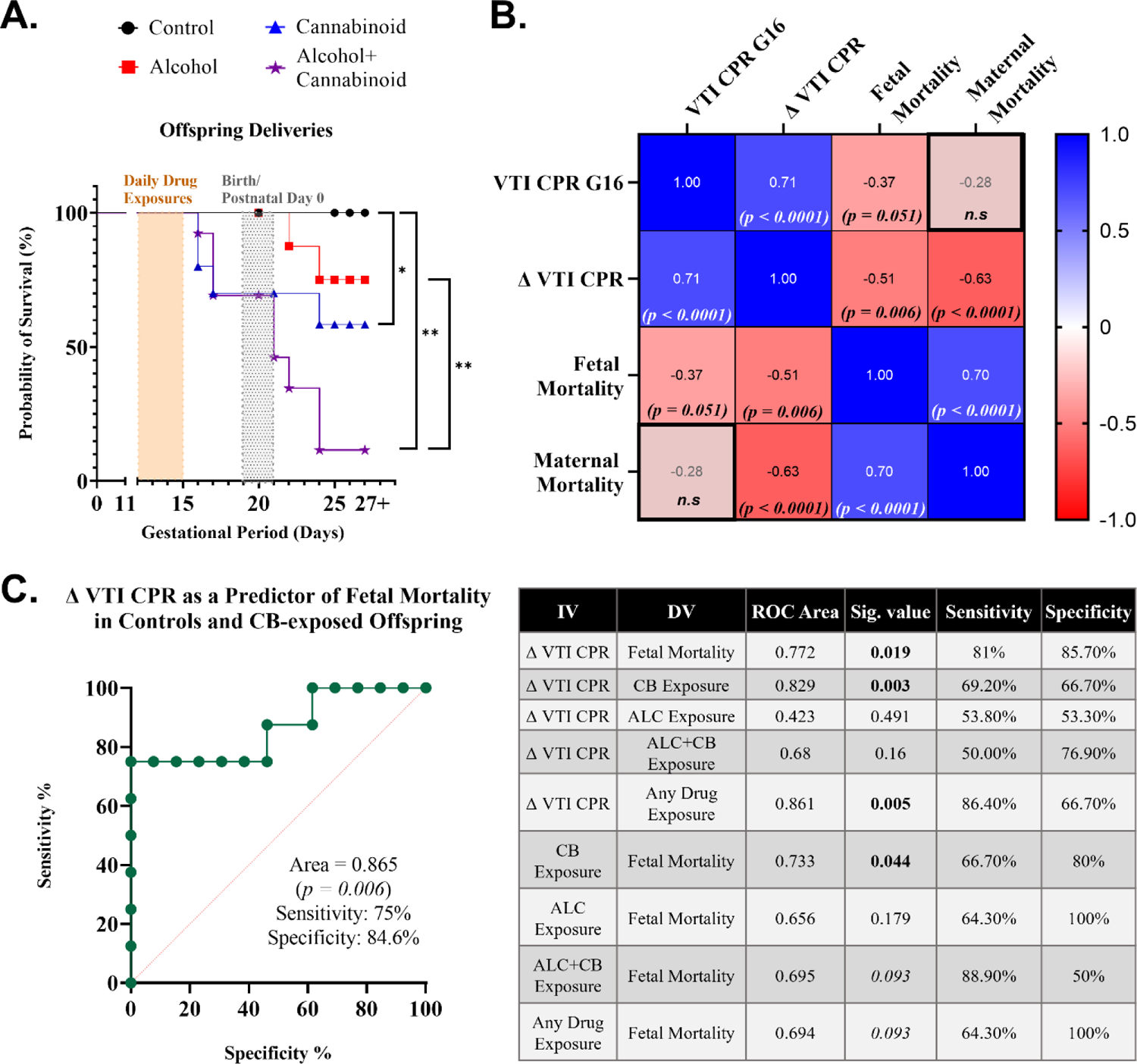
Fetal blood flow metrics as predictors of prior drug exposure and future perinatal mortality. **4A)** Kaplan-Meier analysis of offspring survival rates across drug-exposed pregnancies. **4B)** Pearson’s r correlations between ultrasound measures of cerebroplacental ratios and perinatal mortality among offspring and dams. **4C)** Predictive capacity of cerebroplacental ratios to prior drug exposure or subsequent perinatal offspring mortality. **Abbreviations**: G=Gestational Day, VTI=Velocity-Time Integral, CPR=Cerebroplacental Ratio, ALC=Alcohol, CB=Cannabinoid, ALC+CB=Alcohol+Cannabinoid, N.S=statistically non-significant. **Symbols:** * indicates *p* < 0.05, ** indicates *p* < 0.01.

Compared to Control fetuses, both Cannabinoid (*p*=0.041) and Alcohol+Cannabinoid fetuses (*p*<0.001) were significantly less likely to be born alive. Likelihood of survival for Alcohol+Cannabinoid offspring was significantly worse than Alcohol offspring (*p*=0.004), but not Cannabinoid offspring (*p*=0.115). Maternal survival in the peri-partum period was also marginally compromised in Cannabinoid and Alcohol+Cannabinoid dams [X^2^(3, N=41)=6.385, *p*=0.094; eFig. 4]. In contrast, non-pregnant dams who underwent Cannabinoid and Alcohol+Cannabinoid exposures (*n*=6, data not shown) did not experience subsequent mortality over a 7-month observational period, consistent with prior research showing that CP-55940 is not toxic or lethal to non-pregnant mice at concentrations equal to or greater than those included in our experimental design ^38–40^.

Lower VTI CPR on G16 was associated with increased fetal mortality during the peri-partum period (Fig. 4B). ROC analysis also showed that ΔVTI CPR (from G11-G16) predicted subsequent perinatal mortality (*p*=0.019, area under the curve (AUC), 0.772; sensitivity, 81%; specificity, 85.70%). Δ VTI CPR was also significantly diagnostic of prior exposure to any drug (*p*=0.005; AUC, 0.861; sensitivity, 86.40%; specificity, 66.7%), particularly cannabinoids (Fig. 4C). Finally, gestational exposure to a cannabinoid significantly predicted survival in the perinatal period (*p*=0.044; AUC, 0.733; sensitivity, 66.70%; specificity, 80%). All AUC analyses are detailed in Fig. 4C.

## Discussion

Alcohol is often co-consumed with other psychoactive drugs, but the added risk for pregnancy health and infant birth outcomes due to co-consumption is seldom investigated. SAC in particular requires additional investigation, as existing literature on adult alcohol use disorders indicates that alcohol’s effects may be potentiated by endocannabinoid signaling (reviewed in ^41^). Moreover, two recently published papers in preclinical models indicate that SAC increases teratology ^20,21^ over alcohol or cannabinoid exposures alone. Our study shows that SAC, but not alcohol or cannabinoids alone, decreased gestational weight gain. SAC also augmented maternal blood alcohol concentrations compared to alcohol-exposure alone, a phenomenon that has been observed acutely in human co-users ^42^, as well as rodent offspring with early postnatal exposure^43,44^. As co-use in humans is also associated with higher cannabinoid levels in the bloodstream ^45^, as well as slower cannabinoid metabolism ^46^, it is likely that alcohol and cannabinoids increase each other’s bioavailability. In pregnant individuals, this may indicate that polysubstance use prolongs interactions between consumed substances and a developing fetus.

Whereas all drug exposures decreased fetal cranial blood flow, SAC also reduced internal carotid artery cardiac stroke volume, blood flow quantity, and arterial resistance relative to all other exposures. Moreover, the inhibitory effect of SAC on age-appropriate maturation of middle cerebral and internal carotid blood flow was more severe than following either alcohol or cannabinoid exposures alone. Individually, both alcohol and cannabinoids are known to inhibit fetal cerebral blood flow ^26,34,47–49^, and our data suggest that SAC may augment risk for decreased fetal cranial blood flow. Interestingly, no form of drug exposure inhibited umbilical arterial flow, suggesting a specific vulnerability of fetal cranial circulation.

Both cannabinoid exposure alone and SAC significantly and equivalently decreased fetal CPR, as previously documented for cannabis exposures in human pregnancy ^25^, and in growth-restricted fetuses, we also observed end-diastolic flow similar to observations in human populations ^23^. Reduced CPR suggests a diversion of resources, including nutrients derived from the mother, away from the developing fetal brain. In our study, reduced CPR at the end of the exposure period was a significant predictor of subsequent offspring mortality during the perinatal period. This outcome mirrors findings from one existing early study that reported synergistic increases in fetal reabsorption *in utero* following co-exposure to alcohol and delta-9-tetrahydrocannabinol in rodents ^50^. While cannabinoid exposure by itself has been inconsistently associated with neonatal mortality in human populations ^16,51^, its co-use with alcohol use may confer additional and unrecognized risk for adverse birth outcomes in human populations.

The mechanisms contributing to the synergy between alcohol and cannabinoids are not well understood, although prior research in zebrafish and mice indicates that cannabinoid receptor 1 (CNR1) may be uniquely impacted by SAC ^52,53^, as polysubstance exposure augmented neurobehavioral outcomes and birth defects in exposed offspring in a CNR1-dependent manner. In human populations, the relationship between alcohol exposure and brain CNR1 levels is complex; acute exposure is associated with an increase in CNR1 availability, whereas chronic exposure results in decreased brain CNR1 ^54^. Studies in animal models also generally associate chronic alcohol exposure with inhibition of the endocannabinoid system, including CNR1, though acute exposures produce more variable effects (reviewed in ^41^).

Endocannabinoid signaling plays a critical role in placental growth/viability ^55^ and the dilation ^56^ and growth ^57^ of microvessels. It is possible that the outcomes of SAC are mediated by a complex interplay of activation and inhibition of fetal endocannabinoid signaling across the placenta, blood vessels and brain, resulting in reduced fetal growth and viability. However, further research is required to identify underlying mechanisms, and importantly, to identify possible targets for intervention.

### Limitations

Ultrasound imaging was performed under light isoflurane anesthesia, a necessary step to reduce pain and distress in experimental subjects, but one which complicates interpretation of the effects of drug exposure, although drug-control dams were similarly anesthetized. It is notable, however, that a parallel cohort of SAC-exposed pregnancies which did not include ultrasound imaging also experienced similar mortality during the perinatal period.

## Conclusions

In this carefully controlled preclinical study, ultrasound imaging measures of fetal arterial blood flow were reduced by prenatal alcohol or synthetic cannabinoid exposure, which can disrupt normal embryonic growth and neural development, and polysubstance exposure augmented deficits in cerebral arterial blood flow. These intrauterine metrics subsequently predicted an increased risk of perinatal mortality in cannabinoid-exposed fetuses and their mothers, which may contribute, in part, to reports of greater health risks associated with delivery in marijuana-exposed pregnancies. Given the rising potency and accessibility of cannabis products, including synthetic cannabinoids worldwide, research investigating translational models of prenatal drug exposure, including polysubstance models, are urgently needed to develop informed treatment and care for drug-exposed pregnancies.

## Supporting information

eVideo 1. Control Fetus G16 Waveform

eVideo 2. ALC+CB Fetus G16 MCA Waveform

eVideo 3. Control Fetus G16

eVideo 4. ALC+CB Fetus G16

## Video Legends

**Video 1. Middle cerebral artery blood flow in a Control fetus.** Example ultrasound recording (.avi) of blood flow through the middle cerebral artery in a Control fetus on Gestational Day 16 (post-exposure period).

**Video 2. Middle cerebral artery blood flow in an Alcohol+Cannabinoid fetus.** Example ultrasound recording (.mp4) of absent or reversed end-diastolic blood flow through the middle cerebral artery in an Alcohol+Cannabinoid fetus on Gestational Day 16 (post-exposure period).

**Video 3. Umbilical artery blood flow in a Control fetus.** Example ultrasound recording (.mp4) of blood flow in a Control fetus on Gestational Day 16 (post-exposure period), in which segmented blood flow of the parallel-running umbilical artery and vein (red and blue) can be visualized with Color doppler filters.

**Video 4. Umbilical artery blood flow in an Alcohol+Cannabinoid fetus.** Example ultrasound recording (.mp4) of blood flow in an Alcohol+Cannabinoid fetus on Gestational Day 16 (post-exposure period), in which umbilical arterial and venous blood flow could not be separately resolved in Color doppler mode, as seen in the Control fetus (Video 3). Instead, the presence of spatially overlapping red and blue pixels indicate flow reversal, as documented in human fetuses with a history of cannabis exposure.

