## Supplemental Materials for "Reduced fetal cerebral blood flow following prenatal drug exposure predicts perinatal mortality"

**Online-Only Materials**

**Figures and Figure Legends.**

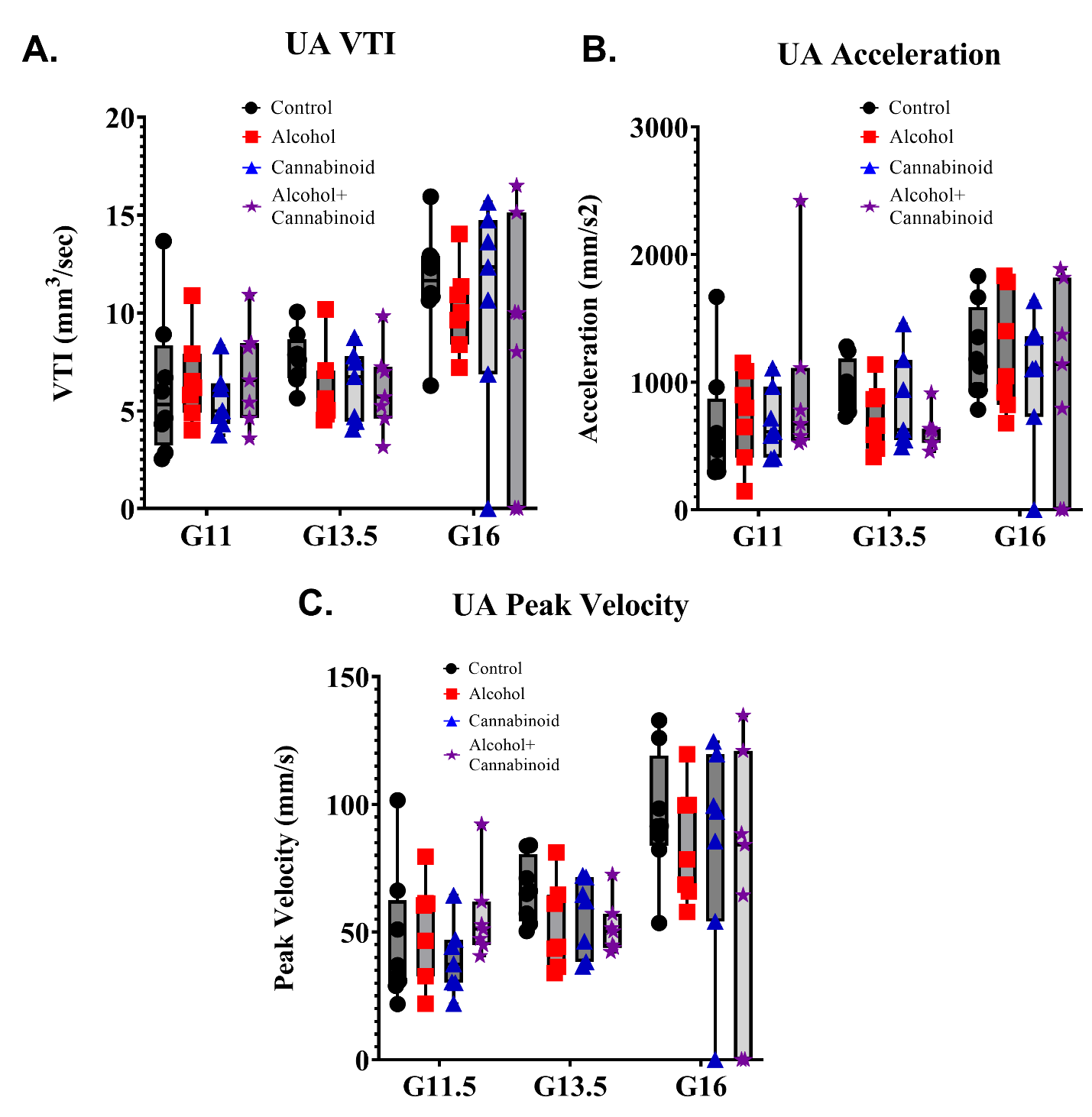

**eFigure 1. Umbilical artery measurements across drug groups and assessment days. 1A)** Assessments of umbilical artery Velocity-Time Integral revealed no significant effect of exposure [*F*(3,25)=0.633, *p*=0.601], nor a significant interaction between gestational day x exposure [*F*(6,50)=0.582, *p*=0.743]. **1B)** Assessments of umbilical artery acceleration revealed no significant effect of exposure [*F*(3,25)=0.179, *p*=0.910], nor a significant interaction between gestational day x exposure [*F*(6,50)= 0.964, *p*=0.459]. **1C)** Assessments of umbilical artery peak systolic velocity revealed no significant main effect of exposure [*F*(3,25)=0.683, *p*=0.571] or interaction between exposure x gestational day [*F*(6,50)=0.776, *p*=0.600]. **Abbreviations**: G=Gestational Day, UA=Umbilical artery, VTI=Velocity-Time Integral.

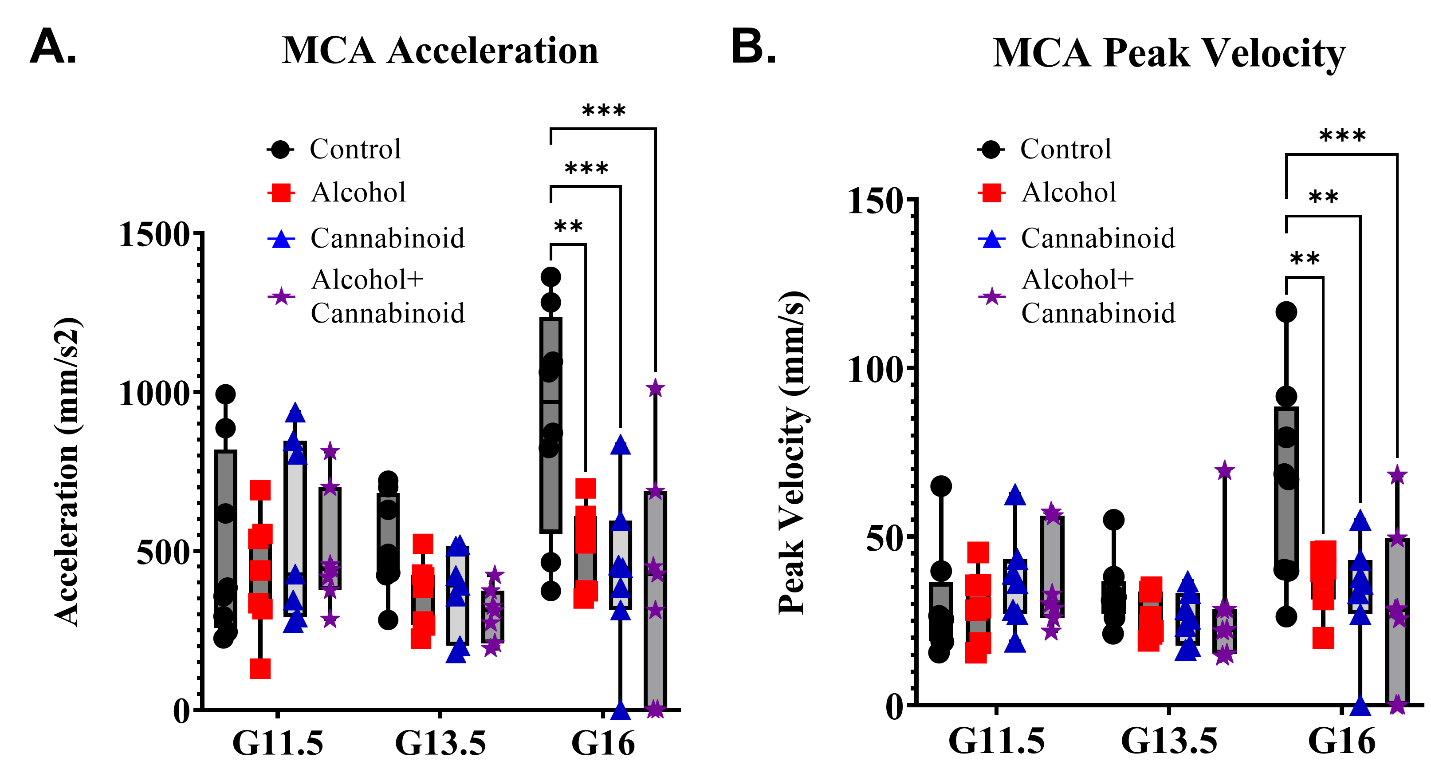

**eFigure 2. Fetal middle cerebral artery measurements of acceleration and peak systolic velocity across drug groups and assessment days. 2A)** Assessments of middle cerebral artery acceleration revealed a significant main effect of exposure [*F*(3,25)=7.297, *p*=0.001] without an interaction between gestational day x exposure (*F*(6,50)=1.899, *p*=0.099). Post-hoc analyses revealed no significant effects of exposure on G11 or G13.5 (eTable 5). On G16, all drug exposures significantly reduced acceleration from Control fetuses: Alcohol (*p*=0.005), Cannabinoid (*p*<0.001), and Alcohol+Cannabinoid (*p*<0.001). Acceleration reductions did not differ between drug-exposed fetuses. **2B)** Assessments of middle cerebral artery peak systolic velocity revealed a significant main effect of exposure [*F*(3,75)=2.837, *p*=0.044] and a significant interaction between gestational day x exposure [*F*(6,75)=3.001, *p*=0.011]. Post-hoc analyses revealed no significant effects of exposure on G11 or G13.5 (eTable 6). On G16, all drug exposures significantly reduced peak systolic velocity compared to Control fetuses: Alcohol (*p*=0.006), Cannabinoid (*p*=0.001), and Alcohol+Cannabinoid (*p*<0.001). Peak systolic velocity reductions did not differ between drug-exposed fetuses. **Abbreviations**: G=Gestational Day, MCA=Middle Cerebral Artery. **Symbols:** ** indicates *p* < 0.01, *** indicates *p* < 0.001.

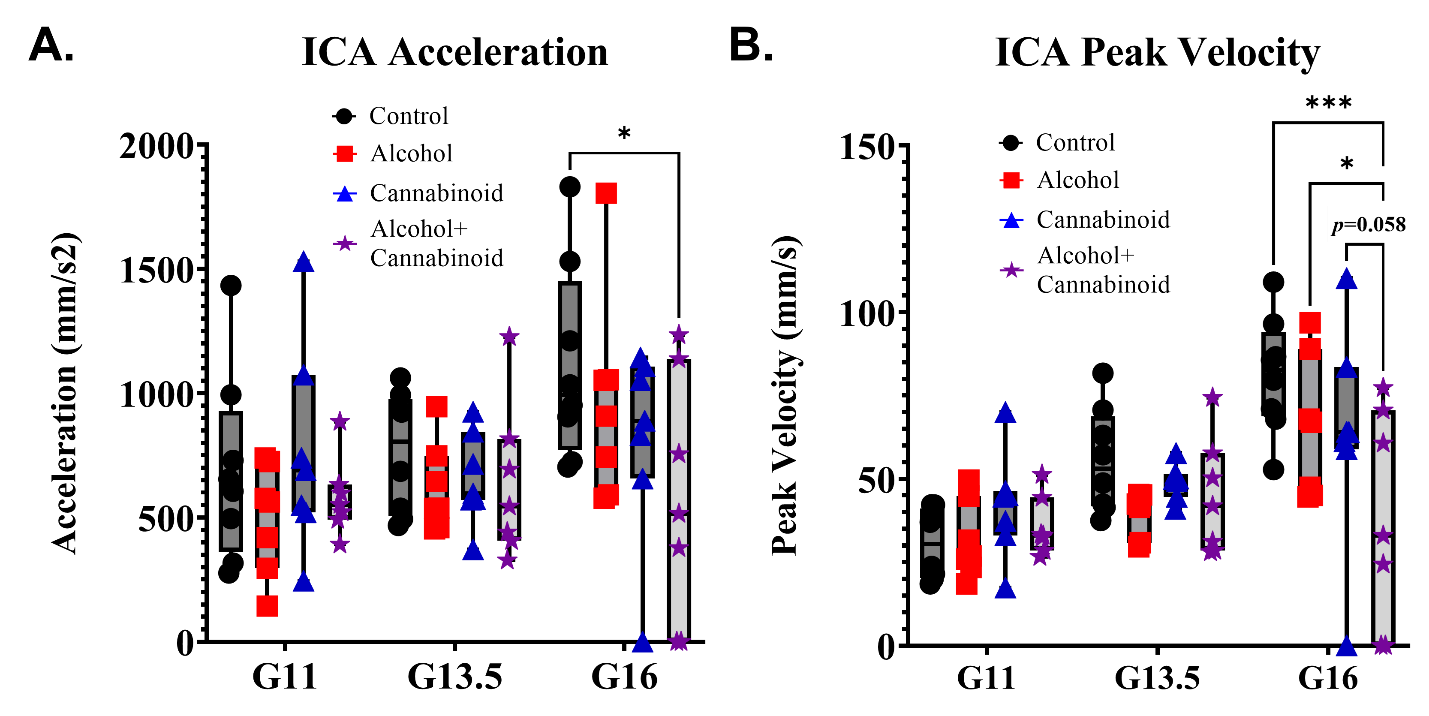

**eFigure 3. Fetal internal carotid artery measurements of acceleration and peak systolic velocity across drug groups and assessment days. 3A)** Assessments of internal carotid artery acceleration revealed a non-significant main effect of exposure [*F*(3,75)=2.269, *p*=0.087], with no interaction between gestational day x exposure (*F*(6,75)=1.152, *p*=0.341). Post-hoc analyses revealed no significant effects of exposure on G11 or G13.5 (eTable 9). On G16, only Alcohol+Cannabinoid fetuses demonstrated reduced acceleration from Control fetuses (*p*=0.015), with no other group differences observed. **3B)** Assessments of internal carotid artery peak systolic velocity revealed a significant main effect of exposure [*F*(3,75)=3.282, *p*=0.025] and a significant interaction between gestational day x exposure (*F*(6,75)=2.651, *p*=0.022). Post-hoc analyses revealed no significant effects of exposure on G11 or G13.5 (eTable 10). On G16, Alcohol+Cannabinoid fetuses demonstrated reduced peak systolic velocity compared to all other exposure groups: Control (*p*<0.001), Alcohol (*p*=0.033) and Cannabinoid (*p*=0.058), with no other group differences observed. **Abbreviations**: G=Gestational Day, ICA=Internal Carotid Artery. **Symbols:** * indicates *p* < 0.05, *** indicates *p* < 0.001. Italicized *p*-value indicates *p* < 0.1.

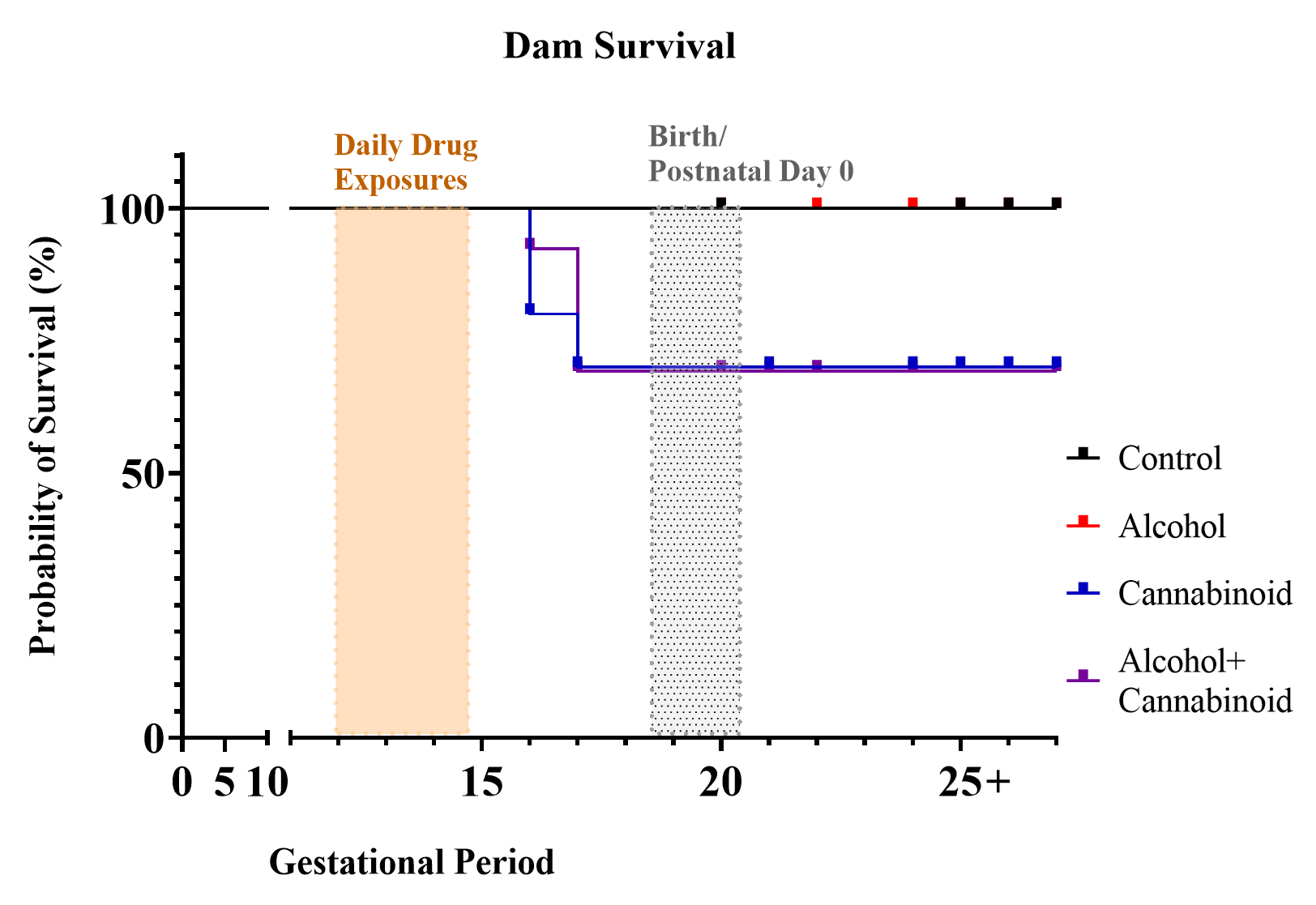

**eFigure 4. Survival rates among dams following drug exposure.** Compromised maternal survival around the time of delivery was observed exclusively in cannabinoid-exposed dams [X^2^(3, N=41)=6.385, *p*=0.094]. Compared to Control dams, maternal mortality increased, although not to statistically significant levels, in Cannabinoid dams (*p*=0.082) and Alcohol+Cannabinoid dams (*p*=0.075), whereas all dams in Control and Alcohol groups survived beyond their estimated date of partition. Importantly, non-pregnant female mice, age-matched to pregnant dams, did not exhibit mortality or other adverse outcomes following an identical drug exposure procedure.
